# Discrete escape responses are generated by neuropeptide-mediated circuit logic

**DOI:** 10.1101/2020.09.22.307033

**Authors:** Bibi Nusreen Imambocus, Annika Wittich, Federico Tenedini, Fangmin Zhou, Chun Hu, Kathrin Sauter, Ednilson Macarenhas Varela, Fabiana Herédia, Andreia P. Casimiro, André Macedo, Philipp Schlegel, Chung-Hui Yang, Irene Miguel-Aliaga, Michael J. Pankratz, Alisson M. Gontijo, Albert Cardona, Peter Soba

## Abstract

Animals display a plethora of escape behaviors when faced with environmental threats. Selection of the appropriate response by the underlying neuronal network is key to maximize chances of survival. We uncovered a somatosensory network in *Drosophila* larvae that encodes two escape behaviors through input-specific neuropeptide action. Sensory neurons required for avoidance of noxious light and escape in response to harsh touch, each converge on discrete domains of the same neuromodulatory hub neurons. These gate harsh touch responses via short Neuropeptide F, but noxious light avoidance via compartmentalized, acute Insulin-like peptide 7 action and cognate Relaxin-family receptor signaling in connected downstream neurons. Peptidergic hub neurons can thus act as central circuit elements for first order processing of converging sensory inputs to gate specific escape responses.

**One Sentence Summary:** Compartment-specific neuropeptide action regulates sensory information processing to elicit discrete escape behavior in *Drosophila* larvae.

---

Animals employ stimulus-specific, optimized strategies to deal with acute threats and noxious stimuli, including escape or avoidance behaviors (*1*–*3*). In the somatosensory system of vertebrates as well as invertebrates, noxious stimuli are sensed by nociceptive neurons and their activation results in acute escape or avoidance (*4*–*7*). A specific noxious stimulus thereby elicits a stereotyped response with high fidelity, e.g. jumping in mice or cork-screw like rolling in Drosophila larvae in response to noxious heat (*6, 8*).

The neuronal networks underlying escape responses show a wide range of complexity from simple reflex circuits to extensive networks regulating threat responses (*8*–*13*). Recent reconstruction of such neuronal networks at the synaptic level in *Drosophila* larvae and neuronal circuit mapping in vertebrates have revealed extensive integration and interaction of circuits mediating distinct responses (*8*–*10, 14*). However, how the appropriate escape behavior is selected in response to a noxious cue is not understood and difficult to deduce from pure anatomical network connectivity, which is only now emerging for more complex nervous systems.

Many neurons across species express neuropeptides (*15*–*18*), which can be released in parallel to small synaptic neurotransmitters and exert modulatory functions (*19*–*22*). Based on their widespread expression and ability to regulate neuronal function, we sought to explore the possibility that neuropeptide signaling plays a role in selective processing of sensory information by the underlying neuronal network to generate appropriate escape behavior.

The *Drosophila* larval somatosensory system presents an opportunity to address this question in an accessible genetic model and by taking advantage of recent advances in circuit reconstruction (*14, 23, 24*). Combined functional studies have revealed a complex network capable of processing different mechanical and noxious stimuli (*14, 25*–*27*) comparable to its vertebrate counterpart (*28*–*30*). At the sensory level, class IV dendritic arborization (C4da) neurons are required for sensing noxious touch and light, which generate two very different escape behaviors (*6, 31, 32*): harsh mechanical touch (mechanonociception) causes corkscrew-like rolling, while exposure to UV or blue light results in reorientation, avoidance and dark preference (termed photonociception). This allowed us to explore how the same circuit can lead to two distinct behavioral outputs depending on the sensory input, and to dissect the underlying neuropeptidergic regulation.

## Results

### Ilp7 releasing neurons are required for photonociception

The dorsal pair insulin-like peptide 7 (Dp7) neurons (*33*) were shown to be required for mechanonociception downstream of mechanosensory (named C2da, C3da and C4da neurons) and 2^nd^ order (A08n) neurons by providing neuropeptide-mediated feedback via the Neuropeptide Y homolog short Neuropeptide F (sNPF, Fig. 1A, A’) (*25*). As Dp7 neurons integrate input from various sensory neurons and have neuromodulatory functions, we reasoned they are likely candidates for computing the discrete C4da neuron-dependent escape behaviors. Similarly to previous studies (*31, 34*), we used a two-choice avoidance assay (darkness vs. white light, 365-600 nm with 6.9-3.3 μW/mm^2^, respectively) to test a potential function of Dp7 neurons in larval photonociception. Inactivation of Dp7 neurons by expressing the inward rectifying potassium channel Kir2.1 with a specific line (*Dp7-LexA* (*25*)) reduced larval light avoidance, suggesting a function in photonociception (Fig. 1B). We next tested whether Dp7 neurons are activated in response to noxious light. Live larvae expressing the calcium indicator GCaMP7s (*35*) in Dp7 neurons were stimulated with a 10 s UV-A light pulse comparable to bright sunlight (360 nm, 60 μW/mm^2^). We found that UV-light exposure gave rise to robust calcium responses in the soma of Dp7 neurons (Fig. 1C), strongly suggesting that Dp7 neurons are part of an innate photonociceptive circuit.

**Fig. 1.**
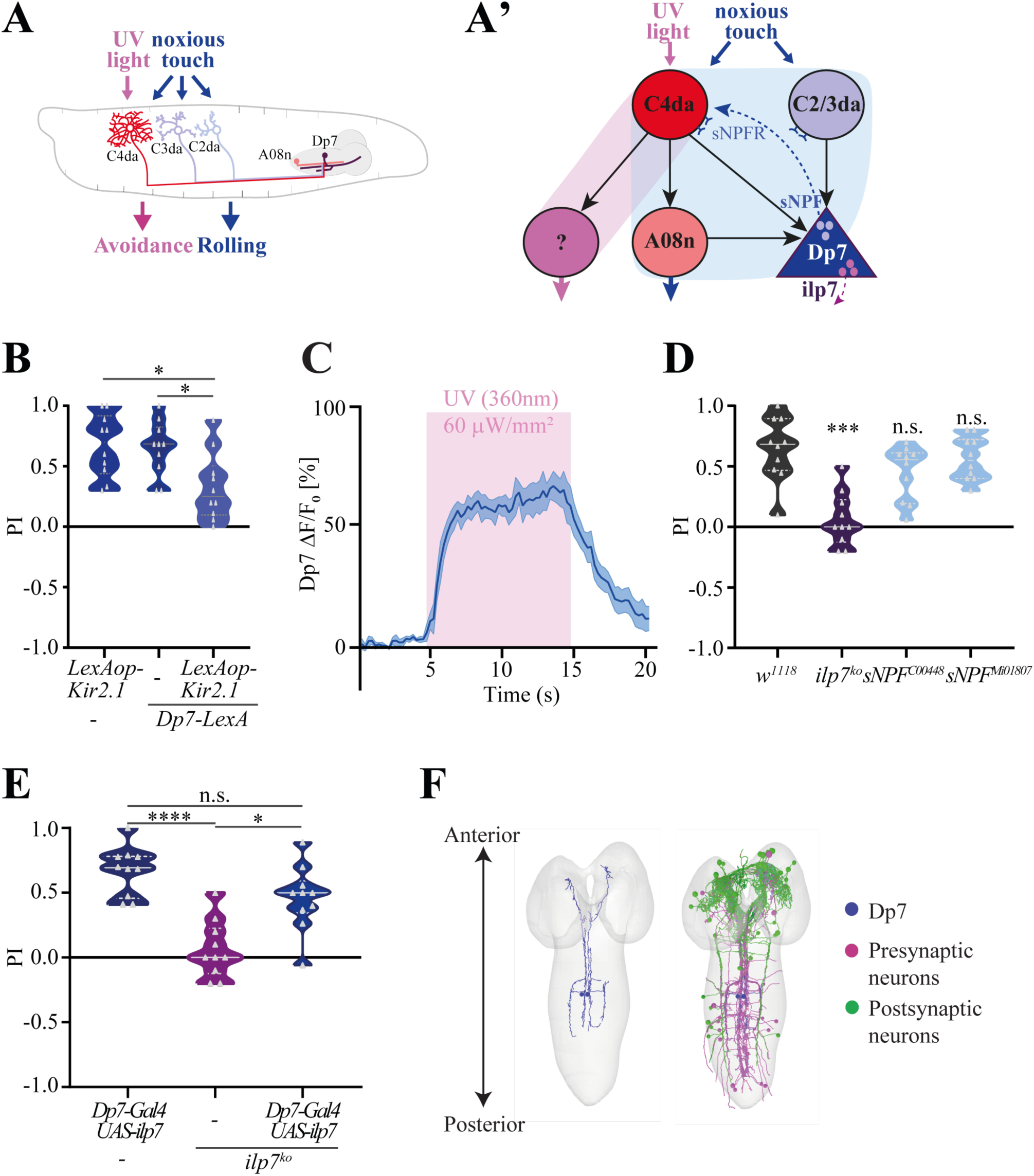
Ilp7-releasing Dp7 neurons are required for photonociception. **A**. Schematic representation of escape behaviors in *Drosophila* larvae. Noxious touch requires C2da, C3da and C4da neurons, while UV light is sensed by C4da neurons which elicits rolling escape or avoidance behavior, respectively. **A’**. For mechanonociception, Dp7 neuron-derived sNPF, but not Ilp7, enables mechanonociceptive rolling through feedback action on C4da neurons to facilitate output to A08n(*25*). **B**. Inactivation of Dp7 neurons using *LexAop-Kir2*.*1* under the control of *Dp7-LexA*, impairs larval photonociception (n=10 trials, ^*^P<0.05, one-way-ANOVA with Tukey’s *post-hoc* test). **C**. UV-A light induces calcium transients in Dp7 neurons (*Ilp7-Gal4>UAS-GCaMP7s*, 365 nm, 60 μW/mm^2^, mean ± s.e.m. indicated by shaded area, n=4). **D**. *Ilp7*^*ko*^, but not *sNPF*, mutant animals showed decreased light-avoidance responses (n=10 trials, ^***^P<0.001, n.s., non significant, one-way-ANOVA with Tukey’s *post-hoc* test). **E**. Dp7-neuron-specific UAS-Ilp7 expression (with *Dp7-Gal4*) in the *Ilp7*^*ko*^ background restores light avoidance (n=10 trials, ^*^P<0.05, ^****^P< 0.0001, n.s., non-significant, one-way-ANOVA with Tukey’s *post-hoc* test). **F**. EM-reconstructed Dp7 neurons and their highest connected synaptic partners. Upstream partners are shown in magenta, downstream partners in green.

To gain a deeper understanding of the modulatory properties of Dp7 neurons in photonociception, we next asked whether the Dp7 neuron-derived neuropeptides sNPF and Insulin-like peptide 7 (Ilp7) are involved in photonociception. Ilp7 affects female egg laying (*36*) and nutrient-dependent tracheal plasticity (*37*), but does not impair mechanonociception (*25*). We analyzed photonociception in *Ilp7* and *sNPF* mutant animals and found that *Ilp7*, but not *sNPF*, is essential (Fig. 1D). To specifically assay the role of Dp7 neuron-derived Ilp7, we rescued Ilp7 expression in *Ilp7* mutant (*Ilp7*^*ko*^) animals using a Dp7-neuron-specific line (*Dp7-Gal4>UAS-ilp7*, Fig. 1E, Fig. S1A). Under these conditions, light avoidance was restored, showing that Dp7 neuron-derived Ilp7 is sufficient for photonociception in *Drosophila* larvae.

To gain more insight into the somatosensory photonociceptive circuit, we first identified the partially reconstructed, but not recognized, Dp7 neurons from the EM brain volume of the first instar larva (*14, 19*). We reconstructed Dp7 neurons and traced all their synaptic partners (Fig. 1F, Fig. S1B-F). The function of Dp7 neurons as a regulatory hub is supported by its connectome as it receives input from several subtypes of sensory neurons (Fig. 1F, Fig. S1E), similar to the hub-and-spoke-like circuit of the *C. elegans* RMG neuron (*38, 39*). Dp7 neurons receive most synaptic input in the ventral nerve cord (VNC) and provide output mostly in the subesophageal zone (SEZ) and brain lobe region (Fig. 1F). Using connectomic analyses, we confirmed connectivity of Dp7 neurons with somatosensory (C2da, C3da, C4da) as well as A08n neurons previously shown at the light microscopic level(*25*) (Fig. S1E). Moreover, we identified a subset of tracheal dendrite (v’td2 (*40*)) neurons as the sensory class with the highest Dp7 neuron connectivity (Fig. S1D). In contrast, the anatomically similar subset of v’td1 neurons was only weakly connected to Dp7 neurons at the connectome level (Fig. S1D,E, see also Fig. 2A). V’td2 neurons are labeled by a reporter line of the putative light sensor Gr28b (*Gr28b*.*c-Gal4*) (*32, 40*) suggesting that, together with previously characterized C4da neurons, they are potential photosensitive neurons on the larval body wall. We confirmed synaptic and functional connectivity between v’td2 and Dp7 neurons using a v’td2-specific Gal4 line (*73B01-Gal4* (*40*)). Synapse-specific GFP reconstitution across synaptic partners (SybGRASP (*41*)) showed that v’td2 neurons form synaptic contacts with Dp7 neurons on their lateral dendrites and the axon (Fig. S2A). Consistently, we detected robust Dp7 neuron calcium responses upon optogenetic activation of v’td2 neurons (Fig. S2B).

**Fig. 2.**
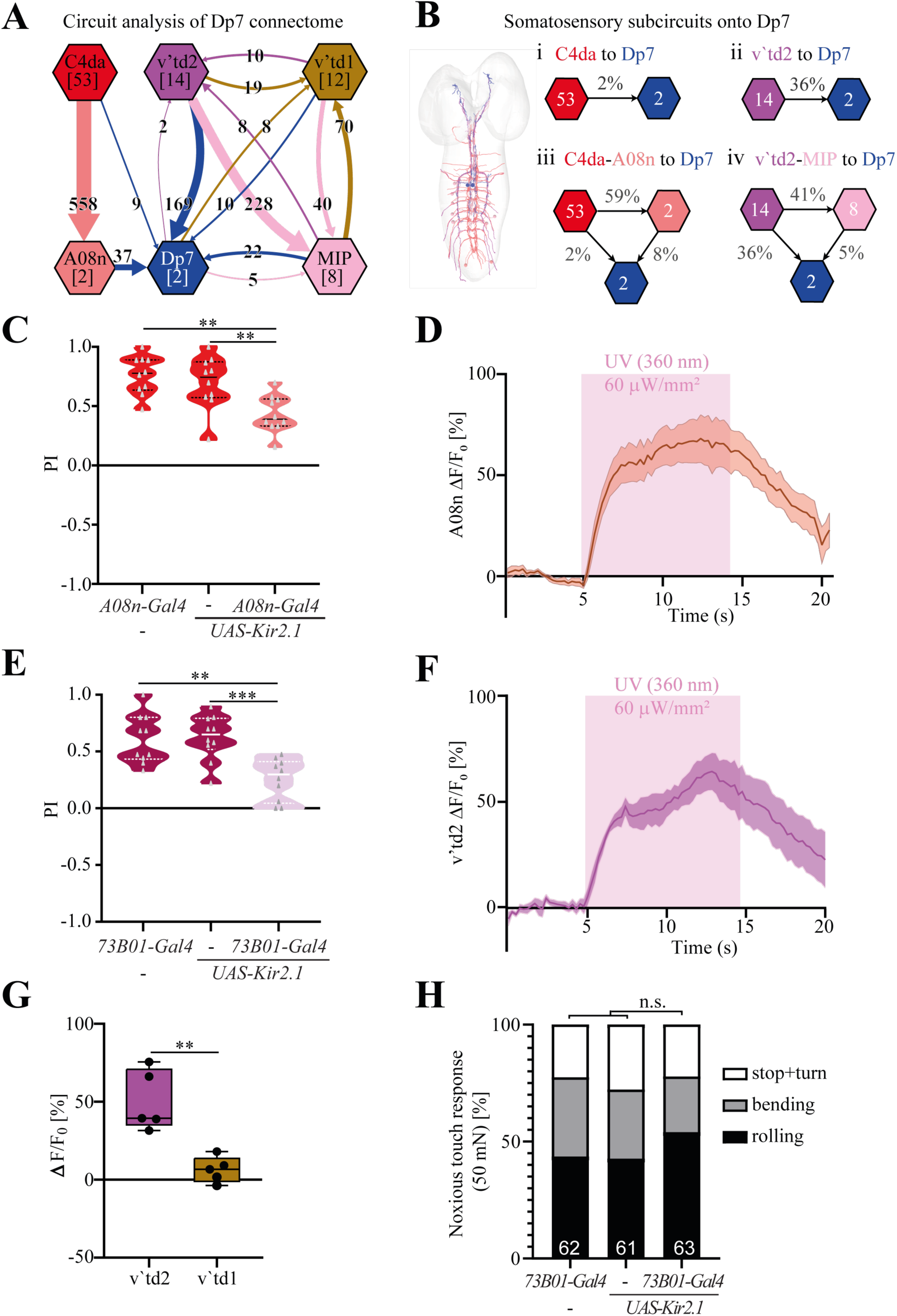
Dp7 integrates noxious light input from multiple somatosensory circuits. **A**. Dp7 neuron presynaptic connectivity analysis showing the highest input from sensory v’td2 neurons. C4da to Dp7 neuron direct connectivity is weak, but additional indirect connections were found via A08n neurons. V’td2 neurons are additionally strongly connected to Dp7 neurons via MIP neurons, while v’td1 neurons display weak connectivity with Dp7 neurons and other circuit elements. Numbers in brackets indicate number of neurons of the respective subtype, numbers on arrows indicate synapses from each neuronal subset forming direct connections. **B**. Inputs onto Dp7 neurons originating from either C4da or v’td2 neurons create 2 direct and 2 indirect subcircuits. Percentages of overall synaptic input of the target cells are shown. **C**. Kir2.1 expression in A08n neurons reduces light avoidance responses (*A08n*-*Gal4>UAS-Kir2*.*1*, n=10 trials/genotype, *P<0.05, **P<0.01, one-way-ANOVA with Tukey’s *post-hoc* test). **D**. Somatic calcium response to UV light in A08n neurons (*82E12-Gal4>UAS-GCaMP6s*, mean ± s.e.m. n=5). **E**. Silencing of v’td2 neurons using *Kir2*.*1* decreases photonociception (*73B01-Gal4>UAS-Kir2*.*1*, n=10 trials, **P<0.01, ***P<0.001, one-way-ANOVA with Tukey’s *post-hoc* test). **F**. UV light-induced calcium transients in v’td2 neurons (*73B01-Gal4>GCaMP6s*, mean ± s.e.m., n=8). **G**. Quantitative comparison of calcium responses (GCaMP6s) of v’td2 and v’td1 neurons to UV light using *R35B01-Gal4*, which labels both subtypes (*Δ*F_max_/F_0_ boxplot, n=5, ^**^P<0.01, unpaired two-tailed *t*-test). **H**. Mechanonociceptive behavior (rolling and bending) is not affected by silencing of v’td2 neurons (*73B01-Gal4>UAS-Kir2*.*1*, n= number of animals as indicated in graph, n.s.=non-significant, *Χ*^2^-test).

### Dp7 neurons integrate noxious light input from multiple somatosensory subcircuits

To uncover the paths through which noxious light information flows in the larval brain, we performed detailed connectomic analyses of the elucidated Dp7 neuron network. We identified four interacting somatosensory circuits converging on Dp7 neurons (Fig. 2A,B). Two of them are direct putative photonociceptive subcircuits (C4da to Dp7 and v’td2 to Dp7 neurons). As the C4da to Dp7 direct synaptic connection is numerically weak and therefore potentially not physiologically meaningful in this behavior, we searched for 2-hop polysynaptic pathways. We identified a strong link via A08n neurons previously shown to receive numerous synaptic inputs from C4da neurons (*23, 25, 42*). In addition, we found that the v’td2 to Dp7 neuron link was strongly interconnected via so far uncharacterized midline projection (MIP) neurons (Extended data Fig. 2C-F).

As C4da neurons respond to UV light and are required for photonociception (*31, 32*), we tested the involvement of A08n neurons as a major downstream output connected to Dp7 neurons. A08n neuron silencing resulted in significantly decreased photonociception (Fig. 2C). In addition, we detected robust calcium transients in A08n neurons in response to UV light (Fig. 2D). Since v’td2 neurons are the major presynaptic partner of Dp7 neurons, we speculated that they are primary sensory neurons transmitting noxious light information to Dp7 neurons. Kir2.1-mediated silencing of v’td2 neurons indeed resulted in significant impairment of photonociception compared to controls (Fig. 2E). We further carried out calcium imaging of v’td2 neurons, which acutely responded to UV light (Fig. 2f). V’td1 sensory neurons, on the other hand, did not show calcium responses to UV stimulation (Fig. 2G), in line with the low connectivity to the Dp7 network (Fig. 2A). These findings show that two sensory subcircuits, C4da-A08n and v’td2 neurons, converge on Dp7 neurons and are involved in UV light sensing and photonociceptive behavior. Unlike for C4da and A08n neurons (*25*), however, v’td2 neuron silencing did not affect mechanonociceptive behavior (Fig. 2H). This shows that v’td2 are specifically linked to UV light avoidance responses, but not mechanonociception.

### A compartmentalized network and acute Ilp7 release transmit photonociceptive information

To identify potential downstream targets of Dp7 neurons, we analyzed the synaptic wiring diagram to identify candidate neurons responding to Ilp7 neuromodulation. Dp7 neuron network analysis revealed numerically weak direct and strong 2-hop synaptic connections of v’td2 via MIP neurons to abdominal Leucokinin (ABLK) neurons (Fig. 3A, Fig. S3A). ABLK neurons have been implicated in blue light-induced rearing behavior through serotonergic input (Okusawa et al., 2014), hinting that they might be involved in photonociception as part of the Dp7 neuron circuit. We inspected the topographical relationship of the mapped neurons and found that v’td2, MIP, and ABLK neurons anatomically converge on the lateral dendrites of Dp7 neurons (Fig. 3B). MIP and v’td2 neurons also overlap with the axonal arbor of Dp7 neurons in the thoracic segments of the larval VNC and SEZ. However, the majority of synapses of the respective postsynaptic partners reside on the Dp7 neuron lateral dendrite (Fig. 3B), suggesting convergence of photonociceptive inputs and outputs. Within this region, Dp7 neurons receive extensive synaptic input from v’td2 neurons, which form concurrent (polyadic) synapses with MIP neurons. MIP neurons, in turn, innervate adjoining ABLK neuron processes (Fig. 3B’). In contrast, the mechanonociceptive circuit comprising C2da, C3da, C4da, and A08n neurons (*25*), of which C4da and A08n also process light information, primarily receives synaptic inputs on the medial dendritic arbor of Dp7 neurons Fig. S3B). This suggests that processing of mechano-and photonociceptive information occurs in distinct Dp7 arbor domains.

**Fig. 3.**
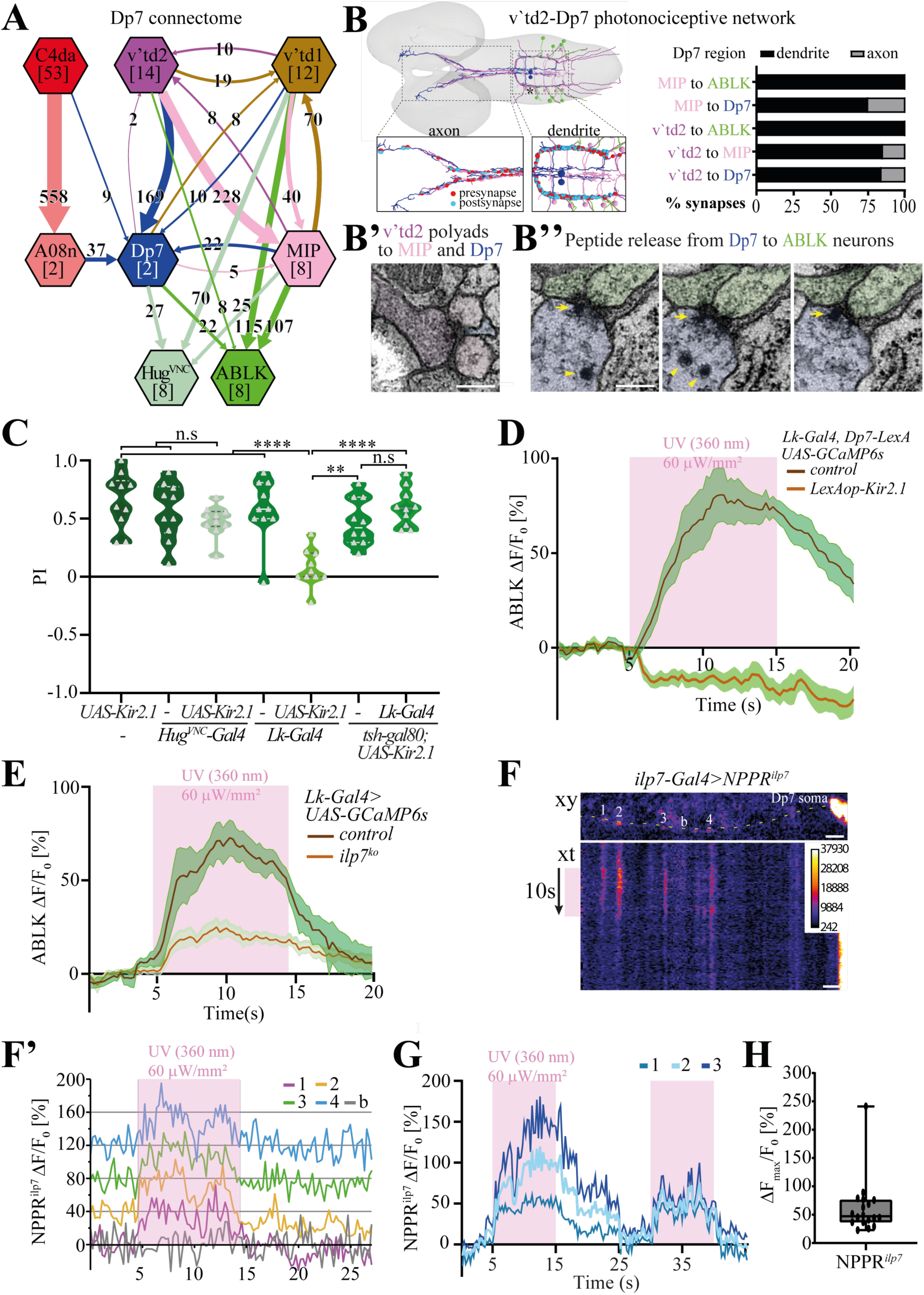
Dp7 neuron activity and acute Ilp7 peptide release is required for photonociceptive information flow to ABLK neurons. **A**. Connectivity graph of Dp7 neurons shows overlapping, but distinct subcircuits. The major outputs of v’td2 neurons are Dp7 and MIP neurons, while v’td1 neurons strongly connect to ABLK and Hugin-VNC neurons. Numbers on arrows indicate synapses from each neuronal subset forming direct connections. **B**. V’td2-Dp7 photonociceptive network. Overview of reconstructed Dp7, v’td2, MIP, and ABLK neuron innervation. Enlarged axon and dendrite regions of Dp7 neurons show local v’td2-Dp7, v’td2-MIP, and MIP-ABLK synapses on the lateral dendrite of Dp7 neurons. Relative synapse numbers in Dp7 dendritic and axonal arbor regions are shown for each partner. **B’**. v’td2 forms polyadic synapses with MIP and Dp7 neurons. Scale bar =200nm. **B’’**. LDCVs (arrowheads) and putative release (indicated by arrow) from Dp7 (blue) to adjacent ABLK neurons (yellow) in consecutive EM sections (shown region indicated by asterisk in Fig. 3b). Scale bar =200nm. **C**. Silencing of LK neurons (*Lk-Gal4*>*UAS-Kir2*.*1*), but not when precluding ABLK expression (*tsh-Gal80, Lk-Gal4>UAS-Kir2*.*1)*, abolishes light avoidance. Silencing Hugin-VNC neurons (*Hug*^*VNC*^*-Gal4>UAS-Kir2*.*1)* does not affect photonociception (n=10 trials/genotype, ****P<0.0001, **P<0.01, n.s., non significant, one-way ANOVA with Tukey’s *post-hoc* test). **D**. ABLK neuron calcium transients evoked by UV light with or without Dp7-neuron silencing (*Dp7-LexA, LexAop-Kir2*.*1*, mean ± s.e.m., n=7). **E**. ABLK neuron calcium transients evoked by UV light in control and *Ilp7*^*ko*^ animals (mean ± s.e.m., n=5). **F**. *NPRR*^*Ilp7*^*-* labeled LDCVs (numbers 1-4, b: background) located along the Dp7 proximal axon. Time series (xt) along the dotted line showing actute evoked *NPRR*^*Ilp7*^ fluorescence increase in response to a 10-s UV-light exposure (360 nm, 60 μW/mm^2^). Scale bars=10 μm. **F’**. Stacked indivdual traces of *NPRR*^*Ilp7*^*-*labeled LDCVs (numbered 1-4) and background (b) shown in **f. g**. Repeated UV-light induced responses of individual *NPRR*^*Ilp7*^ puncta located along the proximal axon or lateral dendrite of Dp7 neurons (from 3 representative experiments). **H**. ΔF_max_/F_0_ boxplot of Dp7 *NPRR*^*Ilp7*^ responses to UV light (n=18 puncta from 6 animals).

Interestingly, the synaptic contact region of v’td2-MIP-ABLK neuron on the lateral dendritic arbor of Dp7 neurons also coincides with Ilp7 neuropeptide localization (Fig. 3B, Fig. S3C), suggesting this could be a site of local peptide release. Analysis of the lateral dendritic arbor of Dp7 neurons in the EM volume revealed putative fusion events of large dense-core vesicles (LDCVs) from Dp7 neurons to neighboring ABLK neurons (Fig. 3B’’, region marked with asterisk in Fig. 3B).

To test whether ABLK neurons are required for photonociception, we performed light avoidance assays using Kir2.1-mediated silencing with a line expressing in Leucokinin (Lk) neurons (*Lk-Gal4*(*43*)), which resulted in significantly decreased photonociception (Fig. 3C). As Lk is expressed in additional neurons in the SEZ and brain lobes, we genetically suppressed expression of Kir2.1 only in ABLK neurons (*tsh-Gal80*). Silencing of the remaining Lk-positive neurons did not result in light avoidance defects suggesting that photonociception is specifically dependent on ABLK neuron function. We also tested Hugin-VNC neuron function in photonociception, which are connected to Dp7 neurons, but receive major sensory input from non-UV responsive v’td1 neurons (Fig. 3A). Consistent with our connectome and functional analysis, we did not detect any significant defects when silencing Hugin-VNC neurons with a specific Gal4 line (*44*) (Fig. 3C), showing that ABLK, but not Hugin-VNC neurons, are specifically involved in photonociception.

We then assayed ABLK neuron responses to UV light using GCaMP6s and found prominent calcium transients upon stimulation (Fig. 3D). To validate that ABLK-neuron responses to UV light depend on Dp7-neuron activity, we silenced Dp7 neurons using Kir2.1 expression. Under these conditions, ABLK-neuron calcium transients were completely absent (Fig. 3D, Fig. S3E). This data is consistent with photonociception requiring information flow from Dp7 to ABLK neurons, either via synaptic transmission or Ilp7 neuropeptide release as suggested by our light microscopic and EM volume analyses. We thus performed calcium imaging in *Ilp7*^*ko*^ animals and detected a 70% decrease in ABLK neuron responses after UV light stimulation (Fig 3E, Fig. S3F). Although ABLK neurons still displayed approx. 30% of the normal responses to UV light in *Ilp7*^*ko*^ animals (Fig. 3E), possibly through synaptic input from v’td2-MIP neurons, this synaptic activity alone was not sufficient to trigger photonociceptive behavior. Altogether, these data indicate that Ilp7 acts on top of the physical photonociceptive connectome to drive noxious light-induced avoidance responses.

Neuromodulation of networks adds a layer of regulation contributing to the computational mechanism required for behavioral action (*15, 17, 22, 45*). Elucidating the spatiotemporal action of neuropeptides is difficult because peptide release can be tonic, independent of synaptic activity, and may act on neurons far away from where they are produced. Based on potential LDCV release from Dp7 to ABLK neurons found in the EM volume at convergence sites of v’td2, MIP, and Dp7 neurons with ABLK neurons, we asked whether Ilp7 release from Dp7 neurons can be acutely induced by UV light stimulation. To visualize peptide release from Dp7 neurons, we generated a release reporter by fusing Ilp7 to GCaMP6s (NPRR^*Ilp7*^), analogously to previously-characterized neuropeptide reporters (*46*). NPRR^*Ilp7*^ expressed in Dp7 neurons localized in a punctate pattern predominantly on lateral dendritic and axonal branches, highly similar to the endogenous pattern of Ilp7 (Fig. S3G). Moreover, the LDCV-specific Synaptotagmin Syt*α* (*47*) colocalized completely with NPRR^*7Ilp7*^ when coexpressed, confirming correct reporter targeting (Fig. S3H). We next imaged NPRR^*Ilp7*^ responses to UV light in Dp7 neurons in live larvae. NPRR^*Ilp7*^ punctae in the proximal axon and lateral dendrite region of Dp7 neurons displayed low baseline fluorescence consistent with low LDCV calcium levels, which should increase upon plasma membrane fusion indicating peptide release. NPRR^*Ilp7*^ fluorescence in LDCVs increased rapidly upon UV-light illumination (Fig. 3F, F’). Repeated UV-light stimulation resulted in consistent NPPR^*Ilp7*^ responses in LDCV puncta (Fig. 3G, H). This data is compatible with acute and rapid peptide release by partial LDCV fusion with the plasma membrane in the millisecond to second range, similarly to described kiss and run-type peptide release upon electrical stimulation (*46, 48*). Imaging of NPPR^Ilp7^ in the Dp7 soma showed similar responses, also suggesting somatic release (Fig. S3I). In contrast, posterior Ilp7-positive neurons, which innervate the gut, did not show an UV-light induced somatic NPPR^Ilp7^ response (Fig. S3I). To further confirm that NPPR^*Ilp7*^ is indeed reporting LDCV fusion with the plasma membrane, we used RNAi to knock down Calcium-dependent secretion activator (Cadps), a conserved protein required for LDCV release, but not biogenesis (*49, 50*). UV-light-induced NPPR^Ilp7^ responses in the Dp7 soma were strongly diminished upon Cadps-RNAi showing that the observed responses are LDCV release-dependent (Fig. S3J). Our data thus show that LDCVs containing Ilp7 are acutely released from Dp7 in response to UV light, thereby acting directly on neighboring ABLK neurons, reminiscent of small molecule neurotransmitter action.

### Neuropeptidergic decoding of photonociceptive circuit responses and behavior

As the photo-and mechanonociceptive circuits overlap extensively at the sensory C4da and Dp7 neuron level, we asked whether Ilp7-dependent output of Dp7 to ABLK neurons is specific for UV light. Kir2.1-mediated silencing of LK neurons, with or without the inclusion of ABLKs, did not significantly impair mechanociceptive escape responses resulting in nocifensive rolling behavior (Fig. 4A). Instead, ABLK neuron silencing mildly facilitated mechanonociceptive behavior, in line with a similar effect described for *Ilp7* deletion (*25*). Moreover, in sharp contrast to UV light stimulation, we did not detect calcium responses in ABLK neurons after mechanonociceptive stimulation (Fig. 4B). Divergence of the mechano-and photo-nocieptive circuits thus occurs downstream of Dp7 neurons through specific Ilp7-mediated action on ABLK neurons.

**Fig. 4.**
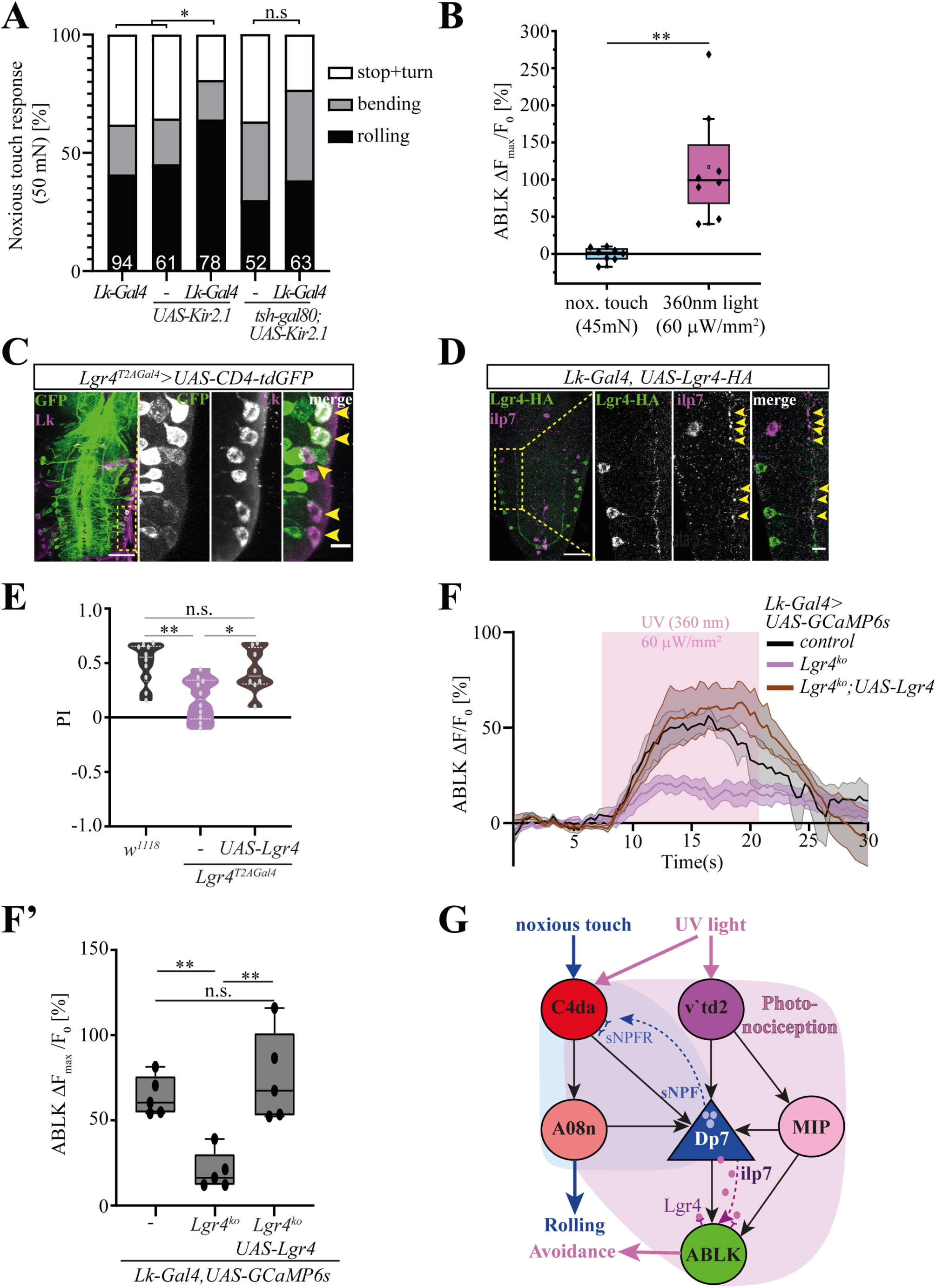
Neuromodulatory decoding of nociceptive escape behaviors. **A**. Mechanonociceptive reponses upon silencing of Lk neurons (*Lk-Gal4 UAS-Kir2*.*1*), with or without ABLK silencing (*Lk-Gal4;tsh-Gal80,UAS-Kir2*.*1*, n=total number of larvae indicated in graphs, n.s.=not significant, ^*^P<0.05, *Χ*^*2*^ test,). **B**. Maximum ABLK neuron reponses (% ΔF_max_/F_0_) to noxious mechanical or UV light stimulations in semi-intact live larval preparations (n=8, unpaired t-test, \*\**P<0*.*01*). **C**. Endogenous Lgr4 reporter expression (*Lgr4*^*T2AGal4*^,*UAS-CD4-tdGFP*) in ABLK neurons detected by colocalized anti-Lk immunostaining. Overview and magnified lateral VNC region (boxed region) with ABLK neuron somata (GFP: green, Lk: magenta). Scale bars=50 μm, 10 μm for enlarged view. **D**. Lgr4-HA localization in ABLK neurons (*Lk-Gal4, UAS-Lgr4-HA*) with anti-Ilp7 immunstaining. Overview and magnified lateral VNC region (boxed region) showing ABLK neuron somata and dendrites with proximity of Lgr4 (green) and Ilp7 (magenta) puncta on the Dp7 neuron lateral arbor. Scale bars=50 μm, 10 μm. **E**. *Lgr4*^*T2AGal4*^ animals display reduced photonociception, which was rescued by *UAS-Lgr4* expression (n=10,10,8 trials/genotype, \**P<0*.*05*, \*\**P<0*.*01*, one-way ANOVA with Tukey’s *post-hoc* test). **F**. GCaMP6s-expressing ABLK neuron responses to UV light in control and *Lgr4*^*ko*^ animals, with and without UAS-Lgr4 expression (*Lk-Gal4*>*GCaMP6s*, n=5 animals/genotype, mean ± s.e.m.). **F’**. Quantitative ΔF_max_/F_0_ box plots of **F** (n=5, \*\**P<0*.*01*, one-way ANOVA,with Tukey’s *post-hoc* test). **G**. Model depicting neural and molecular elements shaping the photonociceptive circuit with specific action of sNPF and Ilp7 on mechano-and photo-nociception, respectively.

While no cognate Ilp7 receptor has been identified so far, the Relaxin-family receptor Lgr4 has coevolved with Ilp7 across arthropod species, suggesting a receptor-ligand relationship (*51*). To determine if Lgr4 is expressed in ABLK neurons and required for photonociception, we first analyzed the expression of a Gal4 reporter incorporated in the endogenous Lgr4 mRNA (*Lgr4*^*T2AGal4*^). We detected Lgr4 reporter signal in ABLK neurons, suggesting it is endogenously expressed (Fig. 4C). We further analyzed the localization of an ABLK-expressed HA-tagged-Lgr4 relative to endogenous Ilp7 neuropeptide using anti-HA and anti-Ilp7 antibodies, respectively. We found that Lgr4 localized along ABLK neuron projections close to Ilp7 punctae on the lateral dendritic branch of Dp7 neurons (Fig. 4D). In addition, we biochemically confirmed Ilp7 and Lgr4 interaction in S2 cells and could isolate specific complexes in co-immunoprecipitation assays showing that Ilp7 and Lgr4 are capable of binding *in vitro* (Fig. S4A,B).

To find out whether Lgr4 is physiologically relevant for photonociception, we tested animals carrying a T2A-Gal4 exon inserted after exon 2, which results in truncation of the endogenous Lgr4 mRNA and loss of *Lgr4* as confirmed by qPCR analysis (*Lgr4*^*T2AGal4*^, Fig. S4C). *Lgr4*^*T2AGal4*^ animals showed significantly reduced photonociception, which could be fully rescued by overexpression of Lgr4 in its endogenous pattern (Fig. 4E). We then asked whether ABLK-neuron responses to UV light require Lgr4. To this end, we imaged calcium responses of ABLK neurons using a confirmed *Lgr4* knockout allele (*Lgr4*^*ko*^ (*52*), Fig. S4D) showing reduced light avoidance as well (Fig. 4F,F’, Fig. S4E). Similarly to *Ilp7*^*ko*^ animals, we detected a three-fold decrease in calcium transients, which could be rescued upon expression of Lgr4 only in LK-positive neurons including ABLKs (Fig 4F,F’). Collectively, these results suggest that Lgr4 acts downstream of Ilp7 in ABLK neurons to promote UV-light responses and photonociceptive behavior.

## Discussion

The emerging *Drosophila* larval connectome of about 10.000 neurons illustrates that sensory networks fan out extensively, adding numerous partners at each subsequent level (*14, 23, 53*). As a result, an unambiguous output path of any given sensory input is often difficult to identify, indicating that physical connection is not a sufficient predictor for function (*17, 54*). Thus central network components with many converging neuronal inputs, like in our case Dp7 neurons, might be general control hubs that gate the activation of specific networks. A similar circuit motif has been identified in *C. elegans*, where the RMG neuron forms a hub and receives spoke-like input from several sensory neurons (*38*). Such convergence of multiple sensory inputs allows integration and modulation of behavioral responses. This might be particularly important in early sensory processing, where peptidergic action can increase the computational power by organizing circuit function to generate alternative behaviors (*17, 22, 55*). Our work revealed that discrete escape pathways are controlled by Dp7 hub neurons through input-specific neuropeptide function. Rolling in response to noxious mechanical touch (*6, 56*) requires feedback signaling from Dp7 neurons via sNPF, but not Ilp7 peptide (*25*). In contrast, photonociceptive avoidance behavior requires Dp7 neuron-derived Ilp7, but not sNPF, and acts via a feedforward mechanism. Circuit-specific neuropeptide action thus generates discrete escape behaviors in this system by creating divergent networks, despite the extensive overlap between mechano-and photonociceptive circuits (Fig. 4G).

Specific compartmentalization of sensory inputs and outputs might further increase the efficiency of network computation at hub neurons through local synaptic domains and neuropeptide release. This is illustrated by the convergence of UV light-responding inputs and outputs with Ilp7 release sites on the DP7 lateral dendritic arbor, which forms a computational unit of the photonociceptive circuit. Discrete functional domains have also been described for *Drosophila* mushroom body Kenyon cells displaying compartmentalized activity, which encodes context-specific functions by dopaminergic modulation (*57*). Thus compartmentalized circuits and neuromodulatory action might be a widespread mechanism to generate context-specific behaviors. Alternative escape behaviors in mice are regulated by competitive and mutually inhibitory circuits of corticotropin-releasing factor and somatostatin-positive neurons in the central amygdala, which mediate conditioned flight or passive freezing, respectively (*10*). While direct involvement of these neuromodulators has not yet been shown, oxytocin release from presynaptic terminals of hypothalamic neurons in the central amygdala attenuates fear responses in mice (*58, 59*), suggesting extensive neuromodulatory regulation of escape and related behaviors across species. Our data further suggests that neuropeptidergic signals can act acutely on the physical neuronal network to promote selective neuronal activation and specific innate behaviors. In general, neuropeptide release has been described to occur upon neuronal activitiy (*20, 60*), yet release probability in neurons *in vitro* is low (*61*) and their action is considered slow and broad (*15, 17*). Thus neuropeptides and cognate GPCRs have well-established roles as slow-acting modulators on targets distant from release sites, e.g., opioid receptor signaling in stress-induced analgesia (*62*), and for regulating long lasting behavioral states including sleep, foraging and social behavior (*39, 63, 64*). Descriptions of acute signaling function in sensory behavior however are rare. Acute neuropeptide release has been detected upon electrical stimulation of *Drosophila* motoneurons (*20, 46*), and Substance P is released from primary nociceptors in mice and required for high stimulus pain responses (*65, 66*). Here, we showed that Ilp7 is acutely released from Dp7 neurons in response to noxious light and acts on downstream ABLK neurons, presumably via its cognate receptor Lgr4 to enable photonociceptive responses and behavior.

Lgr4 belongs to the conserved family of Relaxin receptors, which play important roles in synchronizing growth, hypertension, and feeding (*51, 67*–*71*). Recent work also indicates a role for Relaxin-3 in escape behavior through inhibition of oxytocin-producing neurons in the hypothalamus, a brain region implicated in the modulation of escape responses of vertebrates (*58, 72*). This suggests a conserved role of Relaxin signaling in escape responses.

Based on the widespread expression of neuropeptides and cognate GPCRs across species and various brain regions including in escape circuits (*15*–*18*), neuromodulatory hubs with compartmentalized functions as described here might be a very general circuit motif required for computation of behavior in response to acute sensory stimuli.

## Acknowledgements

We would like to thank M. Petersen and Angela R.M. Dias for excellent technical assistance. C. Wegener, T. Oertner, F. Morellini, J. Parrish and Q. Yuan for comments on the manuscript, B. Ye, B. Hofbauer and C. Wegener for communicating results prior to publication. T. Kazimier for advice and trouble-shooting with Catmaid, L. Herren for supervising Dp7 connectome reconstruction. Stocks obtained from the Bloomington Drosophila Stock Center (NIH P40OD018537) and Vienna Drosophila Resource Center (VDRC, www.vdrc.at) were used in this study. cDNA and cells obtained from the Drosophila Genomics Resource Center (NIH grant 2P40OD010949) were used in this study.

## Author contribution

BNI performed and analyzed most experiments including connectome reconstruction and analysis, photonociceptive behavior and calcium imaging, morphological analysis and wrote the manuscript, AW and FZ performed a subset of the photonociception assays, CH performed and analyzed experiments in semi-intact larval preparations, FMT performed and analyzed mechanonociceptive assays, KS made reagents and performed co-immunoprecipitation experiments, EMV, APC, AM, FH, and AMG developed Lgr4 transgenes, and performed qPCR assays, PSch and MP performed connectome reconstruction and analyses, CHY and IMA developed critical reagents, AC performed and supervised connectome reconstruction, PSo made reagents, contributed to circuit analysis, supervised the work and wrote the manuscript.

## Data and materials availability

The data that support the findings of this study are available from the corresponding author upon reasonable request.

## Supplementary Materials

Materials and Methods

Figures S1-S4

Table S1

References

